## Supplementary figures and images for "Birds multiplex spectral and temporal visual information via retinal On- and Off-channels"

### Supplemental Video 1

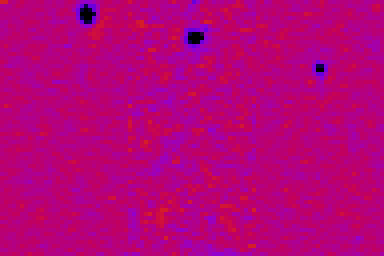
